# Filament-resolved simulations reproduce self-organization of lamellipodia and filopodia

**DOI:** 10.64898/2026.03.15.711798

**Authors:** Masaya Fukui, Yohei Kondo, Nen Saito, Honda Naoki

## Abstract

The dynamic assembly of actin filaments underlies diverse cellular morphologies such as lamellipodia, filopodia, and reticulated networks. However, how filament-scale interactions among actin-binding proteins produce distinct actin architectures remains unclear. We developed a filament-resolved computational model of actin self-organization regulated by the Arp2/3 complex and fascin. Individual F-actin filaments are represented as elastic chains, and their stochastic polymerization, Arp2/3-mediated branching, and fascin-mediated crosslinking and bundling are explicitly modeled. The simulations reproduce three actin architectures observed in minimal reconstitution experiments, including lamellipodia-like branched networks, filopodia-like bundled protrusions, and reticulated meshworks, as a function of Arp2/3 and fascin concentrations. We quantify these regimes using actin density, orientational order, and spikiness, which robustly separate the three morphologies across conditions. To connect filament organization to shape change, we further couple the actin network to membrane deformation using a phase-field formulation. This coupling shows how localized remodeling concentrates load to drive pseudopodial protrusions, whereas highly branched networks distribute stresses and stabilize rounded shapes. The model links molecular interactions to emergent architecture and cell-scale morphodynamics.

## Introduction

Cell morphology is largely determined by the cytoskeleton, especially the network of actin filaments (F-actins) beneath the plasma membrane (Pollard & Borisy, 2003). Through interactions with actin-binding proteins (ABPs), other cytoskeletal components, and the membrane, actin filaments self-organize into higher-order architectures such as lamellipodia and filopodia (Blanchoin et al., 2014; Schaus et al., 2007; Svitkina et al., 2003; Vignjevic et al., 2006; Vinzenz et al., 2012; Welch et al., 1997; Yang & Svitkina, 2011). Directly tracking individual filaments during morphogenesis remains technically challenging, which has limited mechanistic interpretation. Recent advances in live-cell imaging and super-resolution microscopy have begun to resolve actin assembly dynamics and the roles of key regulators including the Arp2/3 complex and formins (Ju et al., 2024; Mueller et al., 2017; Vinzenz et al., 2012; Watanabe & Mitchison, 2002; Yamashiro et al., 2018). Yet a central question remains: how do local filament-scale rules such as branching, crosslinking, and mechanical interactions collectively produce distinct actin architectures at the cell scale, and what sets the transition between lamellipodia-like and filopodia-like organizations (Carlier & Shekhar, 2017; Keren et al., 2008; Tee et al., 2015).

To isolate the essential molecular principles of actin-driven morphogenesis from the complexity of the intracellular environment, in vitro reconstitution systems using a minimal set of cytoskeletal components have been extensively employed. In particular, mixtures of actin monomers, the Arp2/3 complex, fascin, and ATP generate a concentration-dependent phase diagram of F-actin architectures (Haviv et al., 2006; Ideses et al., 2008). High Arp2/3 complex levels yield branched networks reminiscent of lamellipodia, while high fascin favors crosslinked, reticulated meshworks. Notably, at intermediate concentrations, spiny filopodia-like protrusions emerge, suggesting that filopodia can arise from the interplay among F-actins, Arp2/3 complex, and fascin alone. These results imply that key aspects of cellular morphodynamics may be captured by a minimal set of ABPs. However, it remains unclear how filament-scale rules, e.g., branching, crosslinking, and mechanics, collectively generate these distinct regimes and set the boundaries between them.

Many computational models have been developed to study actin-driven morphogenesis, yet a key gap remains: no filament-resolved assembly model has reproduced the concentration-dependent F-actin repertoire observed in minimal reconstitution experiments—lamellipodia-like branched networks, filopodia-like bundled protrusions, and disordered reticulated networks—within a single framework (Haviv et al., 2006; Ideses et al., 2008). Existing approaches broadly fall into two classes.

One class captures cellular morphological dynamics by coupling membrane deformation to intracellular reaction–diffusion systems (Camley et al., 2017; Cao et al., 2019; Marth & Voigt, 2014; Najem & Grant, 2013; Nonaka et al., 2011; Shao et al., 2012), including our previous work reproducing diverse migration modes (Imoto et al., 2021). However, such continuum models typically do not explicitly represent force generation and mechanical interactions at the level of individual filaments. A second class incorporates filament elasticity and network mechanics (Kim et al., 2009; Ma & Berro, 2018; Nedelec & Foethke, 2007; Popov et al., 2016; Weichsel & Schwarz, 2010), but these studies often emphasize local assembly mechanics or specific structures rather than the emergence of cell-scale morphology (Chandrasekaran et al., 2024; X. Chen et al., 2020). Developing a model that captures the emergence of these diverse structures from filament-level interactions would offer crucial insights into the physical principles underlying cellular morphogenesis.

Here we present a filament-resolved computational model in which branching by the Arp2/3 complex and bundling by fascin are sufficient to generate the concentration-dependent phase behavior observed in minimal reconstitution experiments. The simulations reproduce three distinct actin architectures, such as lamellipodia-like branched networks, filopodia-like bundled protrusions, and reticulated meshworks, and quantify them using actin density and filament orientation. Extending the same framework with a phase-field membrane captures reciprocal coupling between cytoskeletal remodeling and membrane deformation, linking filament-scale rules to cell-scale morphodynamics.

## Results

### Filament-resolved mechanochemical model of actin assembly

We developed a mathematical model of self-organization of F-actin assembly to examine how the three different structures, i.e., Network structure, Filopodia-like structure, and Lamellipodia-like structure, emerge. In this model, a single F-actin was addressed as an elastic filament, which was coarse-grained by a one-dimensional connected node. In addition, F-actins were branched by Arp2/3 complex, and cross-linked by fascin (**Fig. 1**). The dynamics of the F-actin assembly can be described as

**Figure 1:**
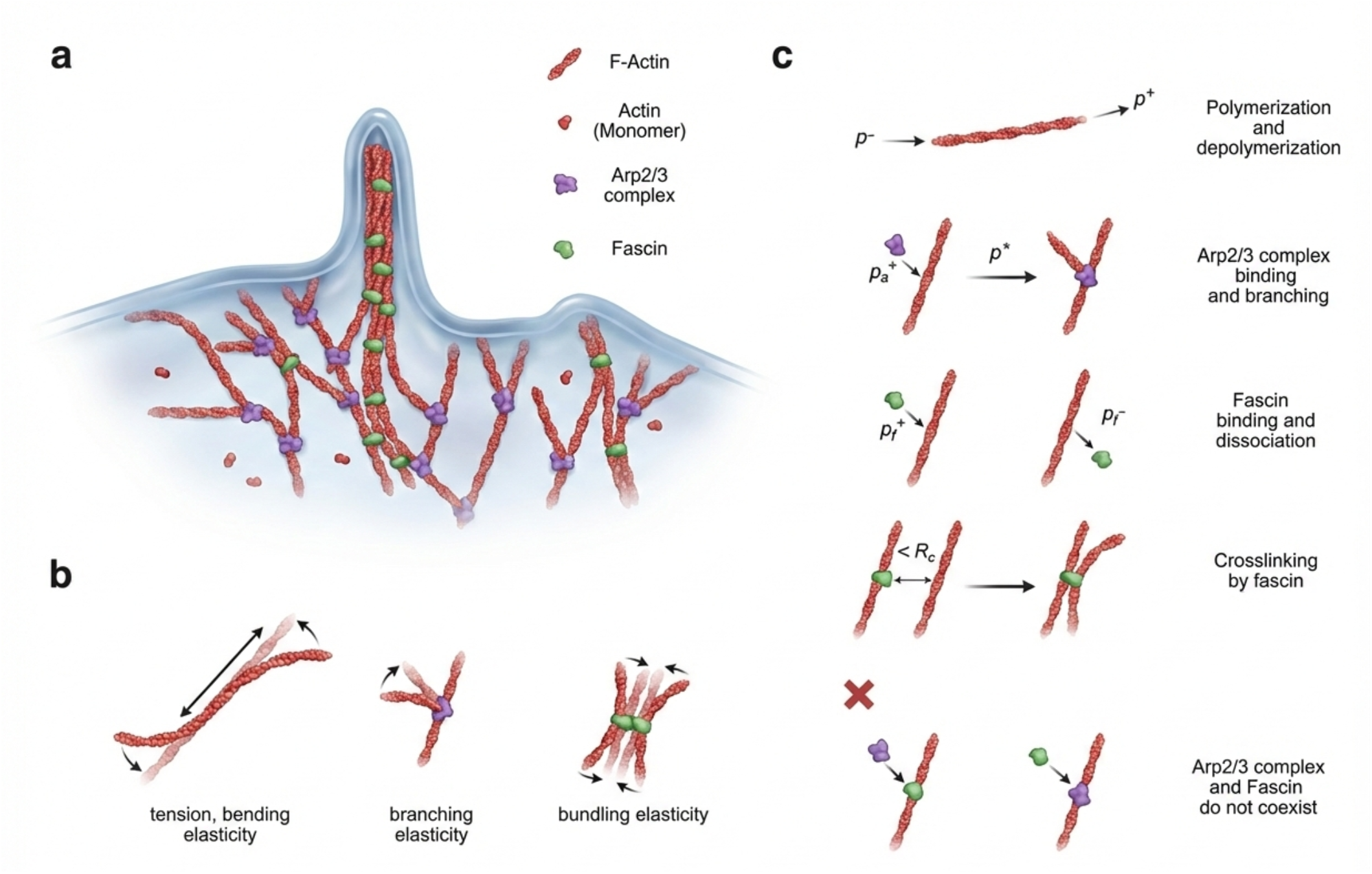
Mathematical model for self-organization of F-actins. **(a)** Our model incorporates key processes governing F-actin dynamics, including actin polymerization and depolymerization, nucleation, branching mediated by the Arp2/3 complex, and bundling facilitated by Fascin. **(b)** The model accounts for the various mechanical properties of the actin filament system, specifically filament tension, bending, bundling, elasticity at branch points maintaining specific angles between two F-actins. **(c)** Chemical reactions underlying F-actin self-organization. The model includes polymerization and depolymerization of actin monomers, binding and branching induced by the Arp2/3 complex, and cross-linking by fascin. It is assumed that the binding of the Arp2/3 complex and fascin to the same filament segment is mutually exclusive.

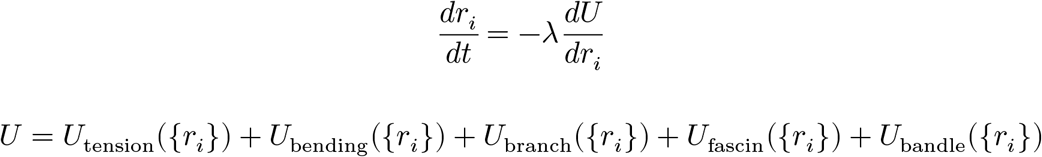

where *r*_*i*_ indicates position of *i*-th node; *U*({*r*_*i*_}) indicates potential energy including tension energy, bending energy, crosslink energy, and branching energy *U*_***_({*r*_*i*_}). *U*_tension_({*r*_*i*_}) represents the generation of tensile force between adjacent nodes, which contributes to maintaining the length of F-actin, whereas *U*_bending_({*r*_*i*_}) represents the generation of bending force, which contributes to maintaining the straightness of F-actin (see Methods in detail).

We performed simulations in 2D to observe the process of actin self-organisation. This means F-actins can cross each other in our simulations.

In the model, F-actins involves several reactions: polymerization/depolymerization, and binding/unbinding with arp2/3 complex and fascin (see Methods in details). Polymerization and depolymerization occur at both ends (Kuhn & Pollard, 2005). The net elongation rate of F-actin at each barbed and pointed end is given by 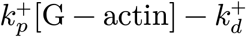 and 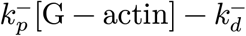, respectively. Note that we considered a situation that the net elongation rates were positive with assumption that G-actins exist abundant in the bulk. Arp2/3 complex associates on F-actin at the rate of *kf*_Arp_[Arp2/3] whereas associated Arp2/3 complex cannot dissociate in the model (Rutkowski & Vavylonis, 2021). Fascin associates to and dissociates from F-actin at the rates of *kf*_Fas_[Fascin] and *kb*_Fas_, respectively. In the model, we assumed that bindings of Arp2/3 complex and fascin were mutually exclusive, namely each actin node can bind either Arp2/3 complex and fascin. Nodes bound by fascin can be connected if they are within the reaction distance.

We assumed that the concentrations are well-mixed. That is, these chemical reaction events can occur with equal probability in all regions in our simulations. We also consider nucleation of F-actin in our simulations. The rate of nucleation is formulated in the same way as for the polymerization of actin. (see Methods in detail)

### Self-organization dynamics of actin networks in filament-resolved simulations

Here, we performed the simulation of the self-organizing process of F-actin networks (**Fig. 2a**). Initially, several F-actin segments were located for seeds of growing F-actin network (**Fig. 2a-1**). After a short interval, the F-actins elongate by polymerization, generate the branched F-actins by Arp2/3 complex and increase the number of total F-actins (**Fig. 2a-2**). When the F-actins elongate and approach each other, the F-actins can be cross-linked by Fascin (**Fig. 2a-3**). During the growth process of the F-actin network, the number of F-acins exponentially increased (**Fig. 2b**) and concentrations of G-actin, Arp2/3 complex, and Fascin monotonically decreased (**Fig. 2c**). To validate this simulation, we confirmed that the potential energy decreases during mechanical relaxation of F-actins, Arp2/3 complex and Fascin (**Fig. 2d**). In this simulation, we visualized spatial locations of Arp2/3 complex and Fascin (**Fig. 2e**), respectively, and spatial distribution of tension and bending forces (**Fig. 2f**), respectively. Thus, our model recapitulated the basic growing dynamics of the self-organized F-actin networks.

**Figure 2:**
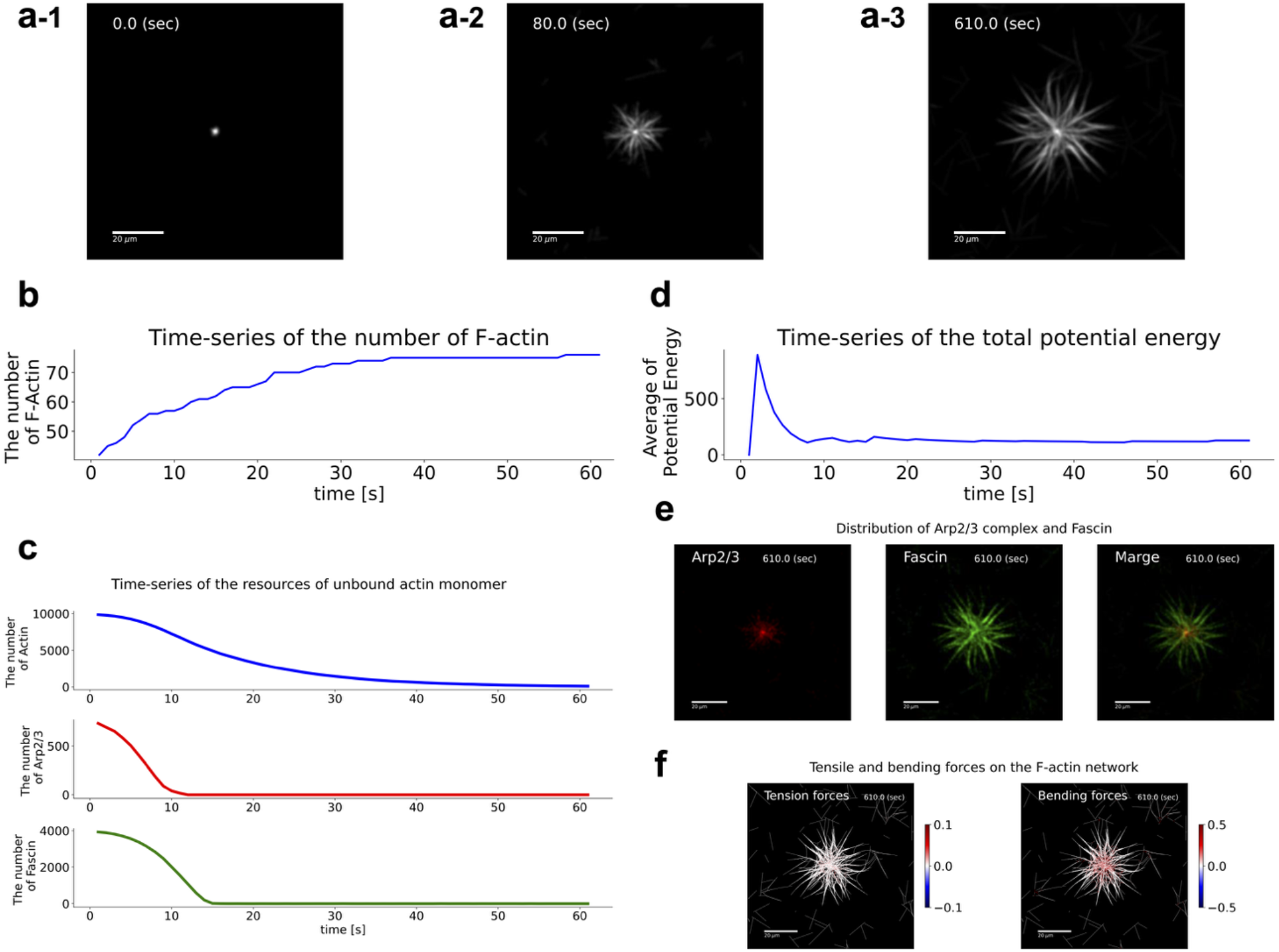
Simulations of self-organizing F-actin networks. **(a)** Growing F-actin network in time. In this simulation, the total number of actin particles is set to 2000, with 800 Arp2/3 complexes and 4000 fascin molecules available as resources. **(b)** Time series of the number of F-actin constituting the structure. **(c)** Time-series of the resources of unbound actin monomer, Arp2/3 complex, and Fascin. **(d)** Time-series of the total potential energy. **(e)** Distribution of Arp2/3 complex and Fascin in filopodia-like structure. **(f)** Visualization of tensile and bending forces on the F-actin network. All images were generated by introducing three new particles between adjacent F-actin particles, rasterizing the F-actin particles to the pixelated image, and then applying a Gaussian filter for a realistic visualization comparable to experimental imaging. All scale bars indicate 20 μm.

### Simulations reproduce three F-actin structures across Arp2/3–fascin conditions

Using the model, we performed simulations, varying concentrations of Arp2/3 complex and Fascin. Then, the model successfully reproduced three distinct types of F-actin structures. With high concentration of Arp2/3 complex and low concentration of Fascin, the F-actin assembly grew into a localized round shape, whose structure corresponds to lamellipodia-like branched networks (**Fig. 3a-1**). This is due to the rapid polymerization from the Arp2/3-induced branching of initially existing F-actins and uniform growth of actin assembly to the surrounding area. On the other hand, with low concentration of Arp2/3 complex and high concentration of Fascin, network structures are generated (**Fig. 3a-3**). In this condition, the localized round shaped-structure did not emerge due to the low concentration of Arp2/3 complex, and then the nucleation of F-actin happened uniformly in space. As a result, F-actins assembled like a network by cross-linking of fascin. In the intermediate condition between two conditions above (i.e., intermediate concentrations of Arp2/3 complex and Fascin), F-actin assembly grew into a localized star shape, which corresponds to a filopodia-like structure (**Fig. 3a-2**). This structure is generated through a process that Arp2/3 complex initially generated the branched-F-actin network, which are then cross-linked by fascin to form bundles. These three emerged structures were consistent with previous in vitro experiments (Haviv et al., 2006; Ideses et al., 2008).

**Figure 3:**
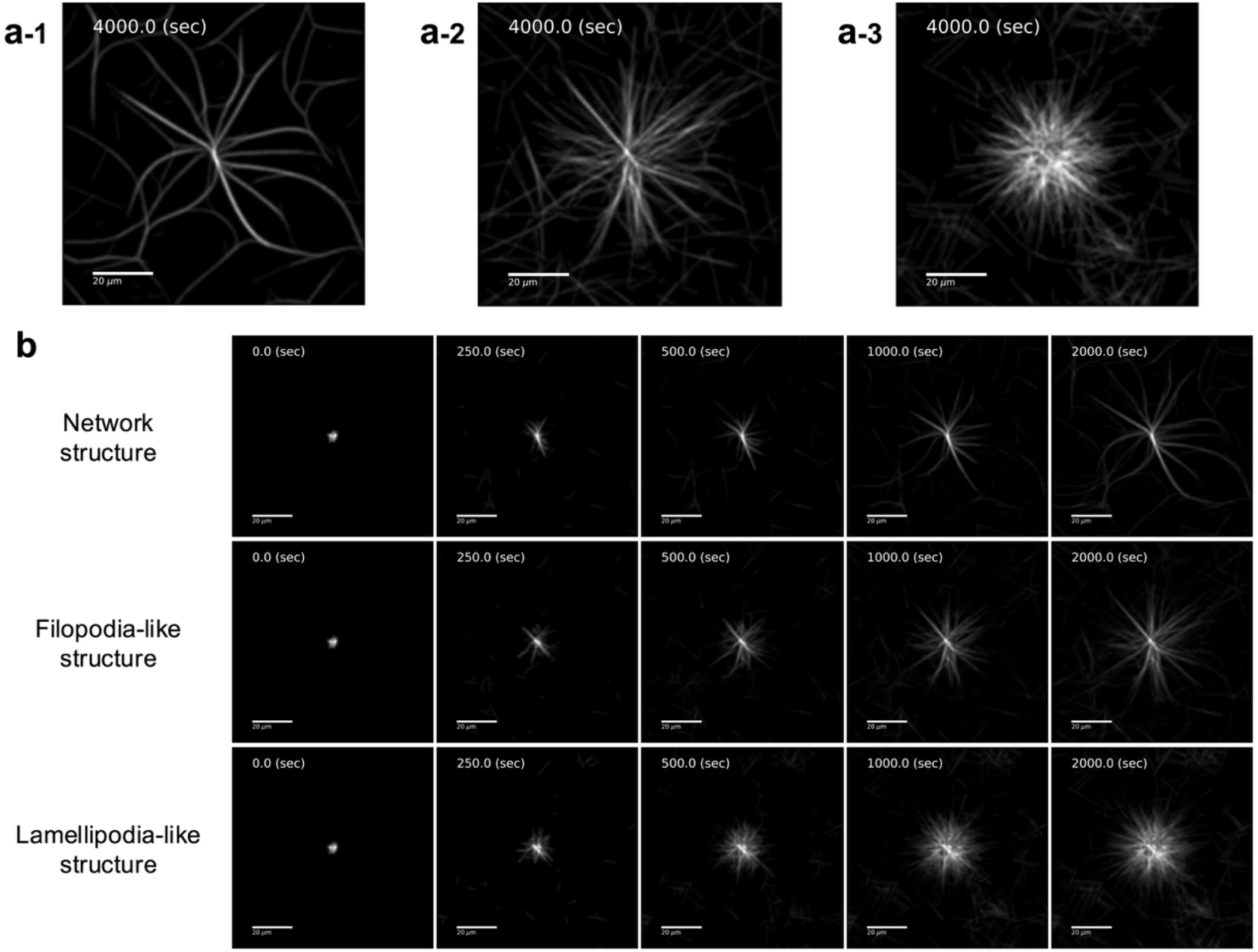
Three types of F-actin structures in simulation. **(a)** Representative simulated image of filopodia-like, lamellipodia-like, and Network structure. The number of actin particles is 2000 in all simulations. Arp2/3 resources are 80 for filopodia-like, 240 for lamellipodia-like, and 640 for network structures, respectively. Fascin resources are 300000, 1000, and 400 for the same structures, respectively. **(b)** Growth process of three types of F-actin structures. All scale bars indicate 20 μm.

### Quantitative evaluations of three types of F-actin structures

We summarized F-actin structures in a phase diagram of Arp2/3 and Fascin concentrations (**Fig. 4a**). To quantitatively evaluate F-actin structures, we characterized them by three different features: F-actin density, orientation order parameter, and degree of spikiness.

**Figure 4:**
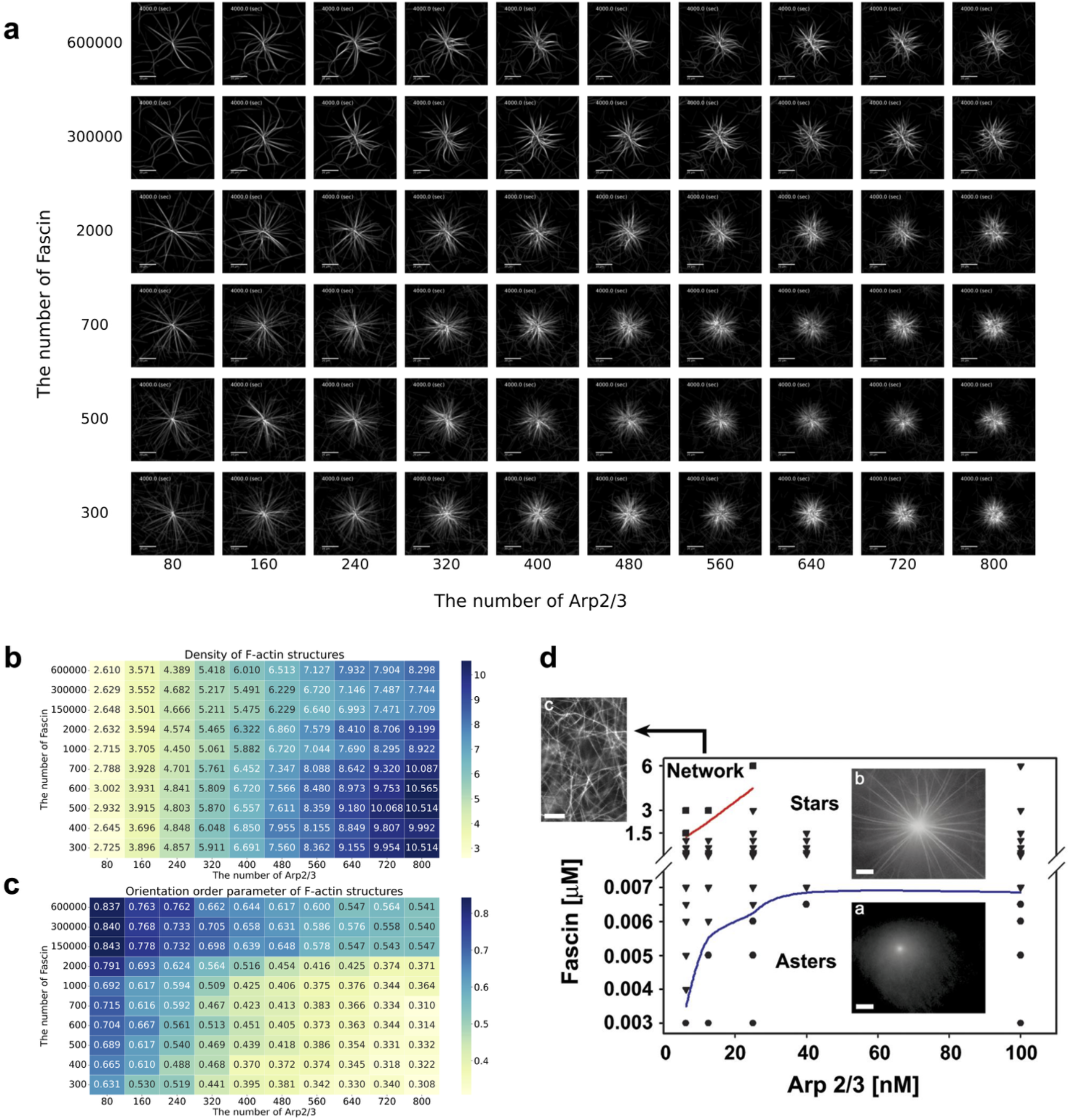
Quantitative evaluations of three types of F-actin structures. **(a)** Phase diagram for the F-actin structures varying concentrations of Fascin and Arp2/3 complex. **(b**,**c)** Heat maps for the density **(b)** the orientation order parameter **(c)** density of F-actin structures. **(d)** Phase diagram of F-actin structures formed with 7 µM G-actin at varying concentrations of fascin and Arp2/3. Structural types are indicated as follows: flamellipodia-like structure (circles), filopodia-like structure (triangles), and network structure (squares). Blue and red lines represent boundaries three types. Adapted from Ideses Y *et al*., *PLOS ONE*, 2008.

F-actin density was calculated by the number of actin nodes within an arbitrary circle, where its radius was selected to ensure that its radius was fixed across all conditions to enable consistent comparison between different structures and concentrations (see Methods). We computed the heatmap of F-actin density in space of the concentrations of Arp2/3 and Fascin (**Fig. 4b**). We found that F-actin density was dominantly determined by the concentration of Arp2/3. Under the conditions of lamellipodia-like structures (i.e., high Arp2/3 and low Fascin) and filopodia-like structures (medium Arp2/3 and high Fascin), the F-actin density tends to be high. On the other hand, under the condition of network structure (i.e., low Arp2/3 and high Fascin), the density became low. Thus, the density feature can separate the lamellipodia/filopodia-like and the network structures, and its boundary was consistent with previous studies.

The orientation order parameter was calculated by the local filament angle (see Methods) (**Fig. 4c**). This order parameter varies depending on the concentration of Fascin and increases continuously in the order of lamellipodia-like, filopodia-like, and network structures. Thus, the orientation order parameter can characterize the three types of structures.

The degree of spikiness is evaluated by the variance of angle of actin nodes, where the positions of actin nodes were represented in the polar coordinate system from the center of F-actin structures. In situations of lamellipodia-like structures (i.e. high Arp2/3, low Fascin) and network structures (i.e. low Arp2/3, low Fascin), the spikiness tends to smaller angular dispersion. Conversely, conditions similar to filopodia-like structures (i.e. medium Arp2/3, low Fascin) correlate with an increased degree of spikiness. Thus, the degree of spikiness distinctly characterizes filopodia-like structures, which is consistent with the phase diagram of F-actin structures observed in previous in vitro study (**Fig. 4d**). Taken together, three types of F-actin structures can be quantitatively separated by three features: F-actin density, orientation order parameter, and the degree of spikiness.

### Membrane coupling links actin architectures to protrusive morphodynamics

So far, we have modeled the mechanism of self-organization of F-actin structures in vitro. However, this model cannot discuss cell morphogenesis because the membrane is not present in this model. Therefore, it is necessary to construct a framework to simulate the dynamics of F-actins surrounded by the cell membrane, rather than simulating only F-actins. Here, we propose a model consisting of two processes: (1) F-actin reorganization within the cell membrane and (2) cell membrane deformation (**Fig 3-a**). As a method to describe the cell membrane, we applied the phase-field method proposed in previous studies (Camley et al., 2017; Cao et al., 2019; Imoto et al., 2021; Marth & Voigt, 2014; Najem & Grant, 2013; Shao et al., 2012; Taniguchi et al., 2013).

We consider the interaction between the cell membrane and intracellular F-actins. We represent the cell membrane using a continuous phase-field field *ϕ* and the F-actin as a chain of discrete sets of points r described earlier. The purpose of this section is to obtain the time evolution equation for some *i*-th vertex *r*_*i*_ and cell *ϕ*. Let *r*_*i*_ be the position of the particle of interest that constitutes the filament. In this study, we represent the filament as a continuous field by considering that this point has a width and can be viewed as a distribution such that 𝒩 (*r*|*r*_*i*_, *σ*_*r*_) in space.

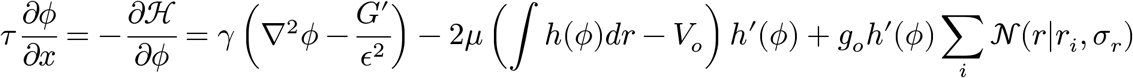

where *G*(*ϕ*) = 18*ϕ*^2^(1 − *ϕ*)^2^ and *h*(*ϕ*) = *ϕ*^2^(3 − 2*ϕ*). The equation of motion of the particle center *r*_*i*_ due to the force exerted by the field *ϕ*(*r*) on the particle center *r*_*i*_ at position *r* is given by

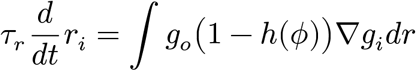

By solving numerically for these time evolutions, the mechanical interactions between the membrane and F-actins can be explicitly treated.

In our simulations, a distinct membrane deformation was identified by varying the Arp2/3 and Fascin concentrations and by adjusting the **membrane tension parameters**. When Arp2/3 concentration was high and Fascin concentration was low, local aggregation of F-actin resulted in a rounded morphology, forming an approximately circular membrane structure (**Fig. 5a-1**). This is due to the branching of F-actin mediated by Arp2/3, which reduces the individual force exerted by each F-actin chain on the membrane (**Fig. 5b**).

**Figure 5:**
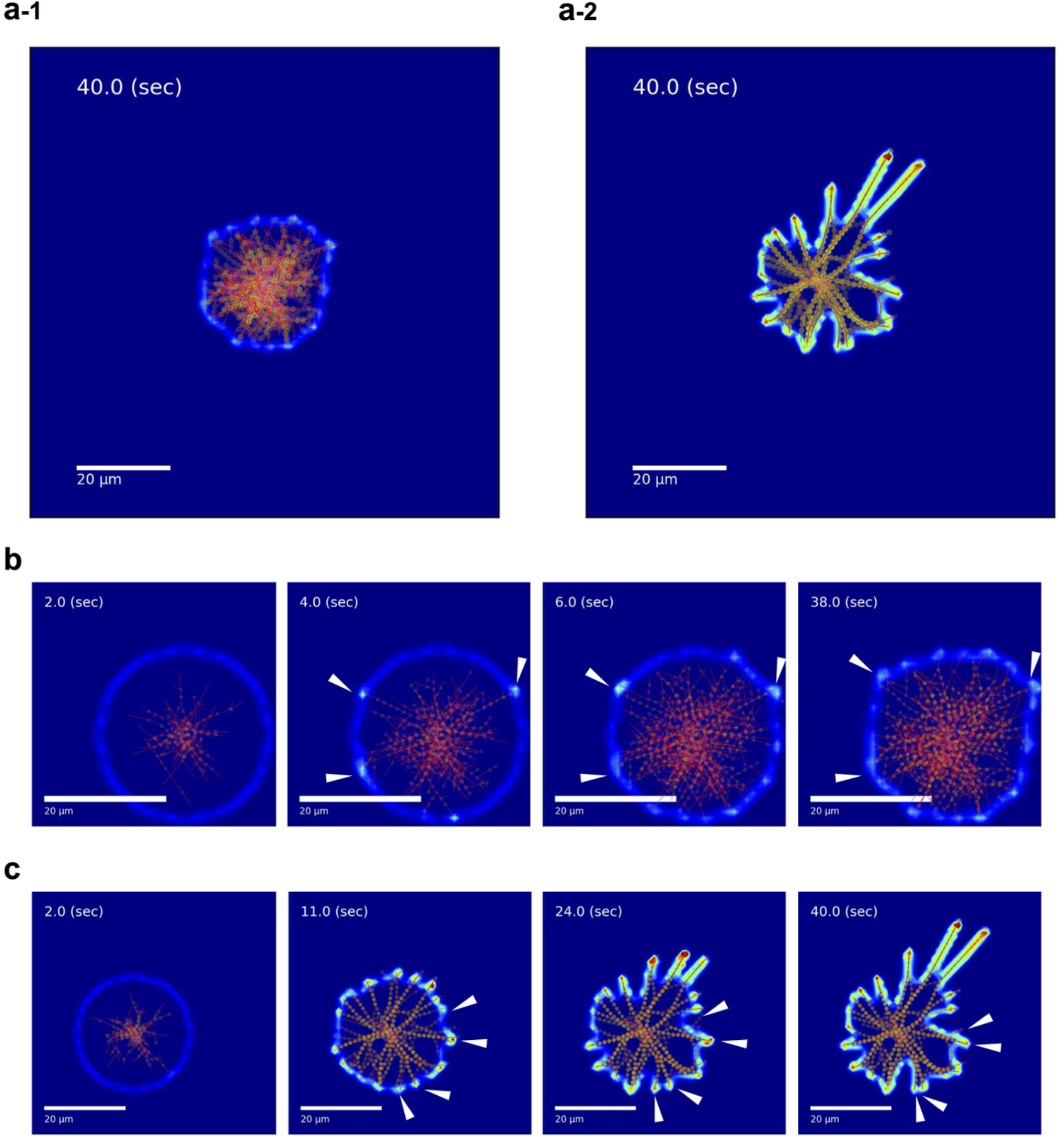
Two types of cell morphology in simulation coupled with membrane dynamics. **(a)** Simulated cell morphology under conditions favoring lamellipodia-like and filopodia-like structures. The heatmap represents the load exerted by F-actin on the cell membrane. Actin is shown in red, Fascin in green, and the Arp2/3 complex in pink. The number of actin particles is 1000 in all simulations. Arp2/3 resources are 80 for filopodia-like and 640 for lamellipodia-like structures, respectively. Fascin resources are 2000 and 640 for the same structures, respectively. **(b)** Snapshots of morphological changes in a cell featuring only lamellipodia. As F-actin presses against the membrane, the load it exerts on the cell membrane gradually reduces. All scale bars represent 20 μm. **(c)** Snapshots of morphological changes in a cell exhibiting pseudopodia. Over time, these pseudopodia are observed to fuse.

In contrast, under conditions of moderate concentrations of Arp2/3 and Fascin, pseudopodia were extended and the membrane was dynamically deformed into complex forms (**Fig. 5a-2**). The bundled and rigid F-actin exerts a <u>strong force</u> to push the membrane forward (**Fig. 5c**). This configuration led to the emergence of complex structures, including filopodia that demonstrated expansion and <u>contraction</u>. Therefore, our mathematical model of cytoskeletal organization within the dynamic membrane is capable of generating both lamellipodia and filopodia formation in a comprehensive manner.

## Discussion

In this study, we developed a computational model that captures the self-organization of F-actins regulated by Arp2/3 and fascin, and demonstrated its ability to reproduce three distinct cytoskeletal structures—lamellipodia-like, filopodia-like, and mesh-like networks—previously observed in vitro. By incorporating filament elasticity, stochastic binding dynamics of regulatory proteins, and membrane deformation using a phase-field approach, our model offers a unified framework to explore how local filament-level interactions give rise to diverse cellular morphologies.

A central contribution of this study is the construction of a physically grounded model that links molecular-scale interactions to the emergence of large-scale F-actin structures. Specifically, we showed that varying the concentrations of Arp2/3 and fascin leads to the formation of different F-actin architectures, consistent with experimental observations. This result highlights the critical role of these two actin-binding proteins in regulating cytoskeletal morphogenesis and demonstrates that complex structural transitions can arise from a minimal set of molecular components. Furthermore, the model integrates filament elasticity and dynamic assembly mechanisms, which are often simplified or omitted in previous models. The ability to simulate lamellipodia-like, filopodia-like, and network-like configurations within a single framework marks a significant step forward in computational modeling of the actin cytoskeleton.

By incorporating membrane dynamics through a phase-field formulation, the model also captures the reciprocal interactions between cytoskeletal assembly and cell shape. Unlike traditional mesh-based methods that impose geometric constraints, the phase-field approach enables seamless coupling between internal filament dynamics and membrane deformation. This methodological choice provides a flexible and extensible platform for simulating cell morphogenesis, with potential for future integration of biochemical gradients, reaction-diffusion systems, or mechanical signaling pathways. Notably, the model remains relatively simple in terms of parameterization, enhancing its interpretability and facilitating further biological applications.

The model is based on biologically plausible assumptions. First, we assume a quasi-two-dimensional system where overlapping filaments can pass through one another. This assumption is supported by previous studies that compared two-dimensional and thin three-dimensional simulations, concluding that two-dimensional modeling sufficiently captures essential dynamics. Second, we assume that each site on a filament can be bound by only one actin-binding protein at a time. This is consistent with biochemical evidence showing mutually exclusive binding among certain regulators, such as between Arp2/3 and capping proteins. Finally, we propose a novel assumption that fascin-mediated bundling occurs preferentially on filaments with limited branching density. While this has yet to be experimentally verified, our simulations suggest that such a mechanism could account for the distinct spatial separation of branched and bundled domains, warranting further experimental investigation.

A number of computational models have been proposed to investigate F-actin dynamics. Early models often neglected filament elasticity (Nonaka et al., 2011), while more recent efforts, such as MEDYAN (Ni & Papoian, 2021; Popov et al., 2016), incorporated mechanical properties like bending and tension, focusing on phenomena such as actin comet tails and contractile rings. However, few models have succeeded in recapitulating the in vitro self-organization of actin structures under minimal conditions. While some models included Arp2/3-mediated branching and fascin-mediated bundling, they did not explore the concentration-dependent transitions among lamellipodia-like, filopodia-like, and network morphologies. On the other hand, phase-field models have been used to simulate cell morphogenesis and membrane deformation (Camley et al., 2017; Cao et al., 2019; Imoto et al., 2021; Marth & Voigt, 2014; Najem & Grant, 2013; Shao et al., 2012). These models typically focused on macroscopic shape dynamics and did not explicitly incorporate filament-level binding rules or cytoskeletal mechanical feedback. Our approach bridges these gaps by integrating microscopic filament dynamics with mesoscopic membrane behavior, thus providing a multiscale modeling framework not previously achieved in this domain.

Despite its strengths, our model has several limitations. For example, the phase-field approach used here does not currently incorporate focal adhesion and actomyosin, which are crucial for linking intracellular actin structures to the extracellular matrix (ECM) and for generating traction forces during migration (Even-Ram & Yamada, 2005; Kuo, 2013). Incorporating such mechanisms would require additional modeling frameworks or hybrid approaches. Additionally, while our current model operates effectively in two dimensions, extension to three-dimensional geometries will be necessary to capture more complex cellular behaviors and morphologies. Finally, the reaction rules and kinetic parameters are based on idealized conditions, and refinements will be necessary to fully align the model with experimental systems. In the future, this modeling framework could be extended to include additional actin regulators (e.g., cofilin, Ena/VASP) (X. J. Chen et al., 2014), signaling pathways, or mechanochemical feedback loops (Hannezo & Heisenberg, 2019). Moreover, the integration of this model with experimental studies may provide a powerful platform for data-driven discovery of cytoskeletal organization principles in cell morphogenesis.

## Methods

### Mechanical model

We developed a mechanical model of the F-actin network represented as a chain of particles connected by springs (Baschnagel et al., 2016) (**Fig. 1**). The positions of the *i*-th particle, *r*_*i*_ = (*x*_*i*_, *y*_*i*_)^T^, evolve following dynamics:

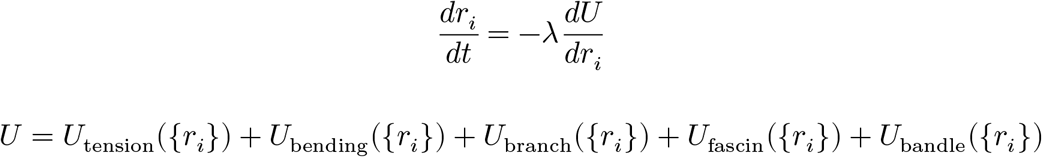

where {*r*} represents the set of *r*_*i*_ (*i* = 1, 2, …); λ is a positive constant; *U*({*r*}) is an energy potential consisting of contributions from tension, bending, branching, and bundling. Each energy term was modelled below:

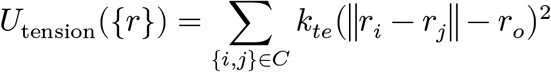

where *C* is the set of connected particle pairs, *k*_*te*_ is the tension stiffness, and *r*_0_ is natural length of the spring (Gittes et al., 1993; Wisanpitayakorn et al., 2022). The bending energy penalizes deviations from a straight configuration:

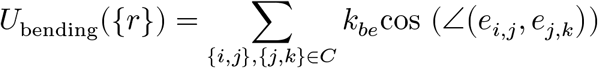

where *k*_*be*_ is the bending stiffness, ∠(*x, y*) represent the angle between unit vectors *x* and *y*, and *e*_*i,j*_ = (*r*_*j*_ − *r*_*i*_)/|*r*_*j*_ − *r*_*i*_|.

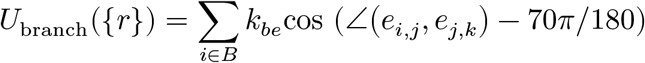

where *B* is the set of particles at the branching point; the *i*-th particle is the branching point connected to the *j*-th and k th participles toward the barbed end direction; *kb*_*r*_ is the bending stiffness at branching points; 70π/180 represents natural angle of 70 degrees which is based on well-established experimental evidence that the Arp2/3 complex nucleates new actin branches at an angle of approximately 70 degrees relative to the mother filament (Amann & Pollard, 2001; Dyche Mullins et al., 1998).

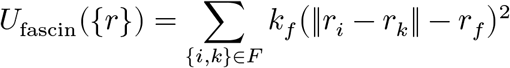

where *F* is the set of particle pairs connected by fascin across different filaments; *kf* is the tension stiffness of fascin; *r*_*f*_ is the natural length of the fascin (Gong et al., 2025; Jansen et al., 2011).

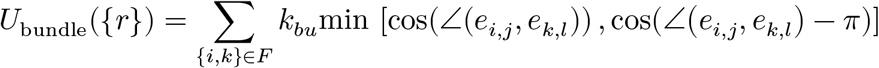

where *k*_*bu*_ is modulus of the fascin-induced force promoting parallel alignment of F-actins.; the *i*-th and *k*-th particles connected to the *j*-th and *l*-th participles within the same filament toward the barbed end direction. Note that min operator represents that if the + ends of the two filaments orient the same direction, the angle formed by the two filaments relaxes to 0 degrees, and if the + ends of the two filaments orient different directions, the angle formed by the two filaments relaxes to 180 degrees. Mechanical model parameters are summarized in **Table 1**.

**Table 1:** Filament-resolved computational model parameters.

| Parameter | Description | Value (*in vitro*) | Value (*in vivo*) |
| --- | --- | --- | --- |
| **Actin filaments** |  |  |  |
|  | The total number of actin particles | 10000 | 2000 |
| $r_o$ | Natural length of the spring [$\mu\text{m}$] | 1.2 | 1.2 |
| $k_{te}$ | Tension stiffness [pN/$\mu\text{m}$] | 100.0 | 100.0 |
| $k_{be}$ | Bending stiffness | 10.0 | 10.0 |
| $k_b^+$ | Barbed end polymerization rate | $2.0 \times 10^{-6}$ | $2.0 \times 10^{-3}$ |
| $k_p^+$ | Pointed end polymerization rate | 0.0 | 0.0 |
| $k_b^-$ | Barbed end depolymerization rate | 0.0 | 0.0 |
| $k_p^-$ | Pointed end depolymerization rate | 0.0 | 0.0 |
| $k_{nu}$ | Nucleation rate | $1.0 \times 10^{-5}$ | $1.0 \times 10^{-3}$ |
| **Arp2/3 complex** |  |  |  |
|  | The total number of Arp2/3 complex | Varied | Varied |
| $k_{br}$ | Bending stiffness at branching points | 10.0 | 10.0 |
| $V_{\text{max},a}$ | Maximum Response | $5.0 \times 10^{-3}$ | 1.0 |
| $K_{m,a}$ | Hill Constant | 160 | 200 |
| $n$ | Hill coefficient | 2 | 2 |
| **Fascin** |  |  |  |
|  | The total number of Fascin | Varied | Varied |
| $l_0$ | Crosslinker spring eq. dist. [$\mu\text{m}$] | 0.1 | 0.1 |
| $k_f$ | Crosslinker spring constant [pN/$\mu\text{m}$] | 50.0 | 50.0 |
| $k_{bu}$ | Crosslinker bending modulus [pN/$\mu\text{m}$] | 10.0 | 10.0 |
| $V_{\text{max},f}$ | Maximum Response | $1.0 \times 10^{-2}$ | 1.0 |
| $K_{m,f}$ | Hill Constant | 480 | 700 |
| $n$ | Hill coefficient | 1 | 1 |
| $k_f^-$ | Dissociation rate of Fascin [1/s] | $4.0 \times 10^{-4}$ | $4.0 \times 10^{-4}$ |

### Reaction model

F-actin undergo stochastic processes such as polymerization, depolymerization, nucleation, and association/dissociation with regulatory proteins including Arp2/3 and Fascin. These dynamics are implemented using probabilistic rules based on the respective reaction rates below.

Polymerization and depolymerization are assumed to occur at both the barbed and pointed ends of filaments. The probabilities of polymerization (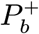 and 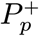) and depolymerization (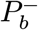 and 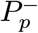)at each end are given by:

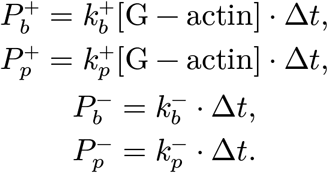

where the superscripts “*b*” and “*p*” denote the plus and minus ends of the filament, respectively; 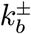 and 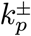 represent the polymerization and depolymerization rate constants; [⋅] denotes molecular concentration; and Δ*t* is the simulation time step. Each polymerization event consumes one actin monomer, while each depolymerization event releases one monomer.

Actin nucleation is modeled as a stochastic process with the following probability:

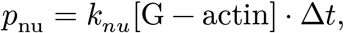

where *k*_*nu*_ is the nucleation rate constant. Each nucleation event consumes two actin monomers and initiates a new filament consisting of two connected particles.

The probability of Arp2/3-mediated branch formation is modeled by Hill-type saturation kinetics:

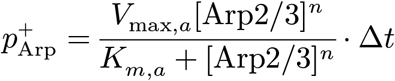

Upon binding, the Arp2/3 complex remains stably associated with the mother filament and nucleates a daughter filament at an angle of 70° relative to the axis of the mother filament. Branching is restricted to a two-dimensional plane, with the daughter filament extending at either +70° or −70° with equal probability. The occurrence of a branching event is determined stochastically according to the probability 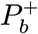 and 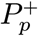.

The probabilities of Fascin binding and unbinding is also modeled using Hill kinetics:

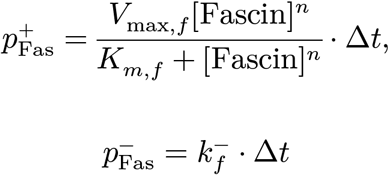

Fascin and Arp2/3 compete for the same binding site on each actin particle, allowing only one of them to bind at a given time (**Fig. 2c**). Upon binding, fascin crosslinks the nearest actin particle from a neighboring filament located within a predefined interaction distance (0.1 μm(100nm)). Fascin dissociation is modeled as a stochastic event that occurs with a constant probability 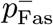 per time step. Reaction model parameters are summarized in **Table 1**.

### Simulation protocol

To achieve high computational efficiency, all simulations were implemented in the C programming language. At each time step, key physical quantities—including particle positions, binding states, filament connectivity, and membrane morphology—were recorded. Post-processing and data visualization were conducted using custom Python scripts, allowing flexible analysis of filament dynamics and structural evolution.

Simulations were performed in a two-dimensional square domain with periodic boundary conditions. The system was initialized with 40 short F-actin fibers, each consisting of two connected particles generated via nucleation. These filaments were randomly placed within a circular region centered in the domain, with a radius of 2μm. The concentrations of free G-actin, Arp2/3 complex, and Fascin were assumed to be spatially uniform throughout the simulation, under the assumption that their diffusion is sufficiently fast relative to the timescale of filament dynamics.

Each actin particle was represented as a C-language structure containing the following attributes: a particle index, spatial coordinates, binding states with regulatory proteins (e.g., Arp2/3, Fascin), indices of neighboring connected particles, and crosslinking information mediated by Fascin. In addition, each particle could form connections at up to three distinct sites: (i) to a neighboring particle in the barbed-end (+) direction, (ii) to a neighboring particle in the pointed-end (−) direction, and (iii) to an additional neighboring particle in the barbed-end (+) direction in the case of branching. These connection states were stored as a three-element binary vector (0 or 1), indicating the presence or absence of a connection at each corresponding site.

Biochemical events such as nucleation, polymerization, depolymerization, and binding/dissociation of Arp2/3 and Fascin were modeled as stochastic processes based on reaction probabilities. Upon nucleation, two actin monomers were consumed; polymerization and depolymerization altered the amount of free G-actin accordingly. Similarly, when Arp2/3 or Fascin bound to or dissociated from filaments, the corresponding molecule counts were updated to reflect these changes. In the simulation, concentrations of all diffusive components were internally represented as discrete molecular copy numbers. When multiple events—such as polymerization, Arp2/3 binding, and Fascin binding—could potentially occur at the same actin particle during a given time step, the actual outcome was sampled based on the relative probabilities of all possible events.

### Feature Quantification for Structural Classification

The simulated F-actin structures were quantitatively evaluated by two feature quantities: density and orientation order parameter. These metrics were used to construct a phase diagram (**Fig. 5**).

Density is defined as the local concentration of F-actins around the center. It is defined as:

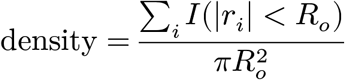

where *r*_*i*_ denotes the position vector of *i*-th actin particle relative to the center of the domain, and *I*(|*r*_*i*_| < *R*_*o*_) is an indicator function that takes the value 1 if |*r*_*i*_| < *R*_o_, and 0 otherwise. Here, *R*_*o*_ defines the radius of the region of interest.

The orientation order parameter (Mottram & Newton, 2014; Steinhardt et al., 1983) was calculated to quantify the angular correlation among neighboring F-actin. This parameter reflects the degree of filament alignment, with higher values indicating more ordered, bundled structures such as filopodia. To compute this value, the region of interest was divided into a fine lattice grid. For each lattice site, we calculated the angular correlation among F-actins present within that site, and then averaged the values across all lattice sites to obtain the global order parameter *S* as

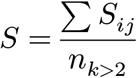

where *n*_*k*>2_ indicates the number of lattice sites containing two or more filaments. *S*_*l*_ represents the angular correlation among neighboring F-actins within the *l*-th lattice as

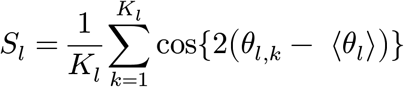

where *k* is the index of the F-actin, *K*_*l*_ is the total number of filaments within the *l*-th lattice site, *θ*_*l,k*_ is the orientation angle of the *k*-th filament, and < *θ*_*l*_ > is the average angle of all filaments within that lattice site (i.e., 〈*θ*_*l*_〉 = (1/2) ⋅ atan(∑_*k*_ sin 2*θ*_*l,k*_, ∑_*k*_ cos 2*θ*_*l,k*_)). Note that the orientation angle *θ* represents the direction of F-actin regardless of polarity, i.e., filaments that are parallel but point in opposite directions are regarded as having the same angle.

### Phase-field models

The total energy of the system is given by the following Hamiltonian:

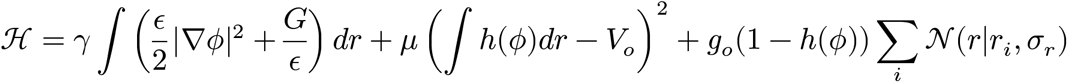

In this formulation, the first term represents the membrane surface tension, where γ is the surface tension coefficient and ϵ determines the characteristic thickness of the interface. The potential *G*(*ϕ*) = 18*ϕ*^2^(1 − *ϕ*)^2^ is a Landau-type double-well potential that stabilizes the interior and exterior regions of the cell. The second term imposes an area constraint using a smooth indicator function *h*(*ϕ*) = *ϕ*^2^(3 − 2*ϕ*), with μ as the strength of the constraint and *V*_*o*_ as the target area. The third term represents a repulsive interaction between the membrane and F-actins, where *i*-th particle of each filament at position *r*_*i*_ is modeled by a Gaussian kernel 𝒩 (*r*|*r*_*i*_, *σ*_*r*_) with mean of *r*_*i*_ and variance of *σ*_*r*_. The coefficient *g*_*o*_ determines the strength of this interaction.

The membrane field *ϕ* evolves according to a reaction–diffusion-type equation obtained from the variational derivative of the total energy functional ℋ with respect to *ϕ*:

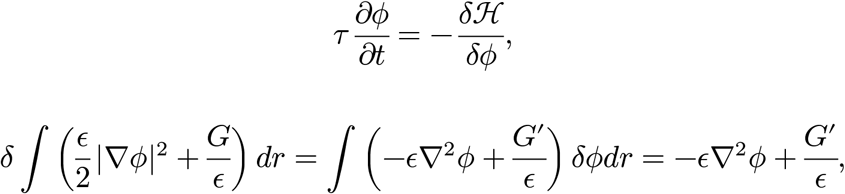

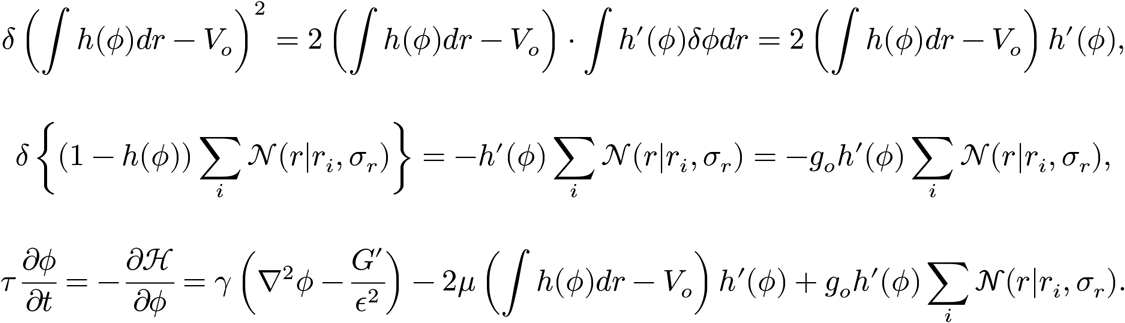

This equation incorporates three physical effects. The first term corresponds to the minimization of membrane curvature via surface tension. The second term restores deviations from the target area, maintaining approximate volume conservation. The third term introduces mechanical feedback from the cytoskeleton by representing how the presence of filament particles locally deforms the membrane. Taken together, these terms define a mechanochemical membrane model that dynamically adapts to intracellular cytoskeletal activity.

### Interaction between F-actin and membrane

The influence of membrane deformation on actin dynamics was incorporated by introducing feedback from the membrane field *ϕ*(*r, t*) to individual F-actins. Each F-actin is modeled as a chain of discrete particles *r*_*i*_, which interact with the membrane through the same Gaussian kernel *g*(*r* − *r*_*i*_) as defined in the energy functional.

The force exerted by the membrane on each actin particle is derived by taking the functional derivative of the total energy ℋ with respect to the particle position. This results in the following equation of motion for particle *r*_*i*_:

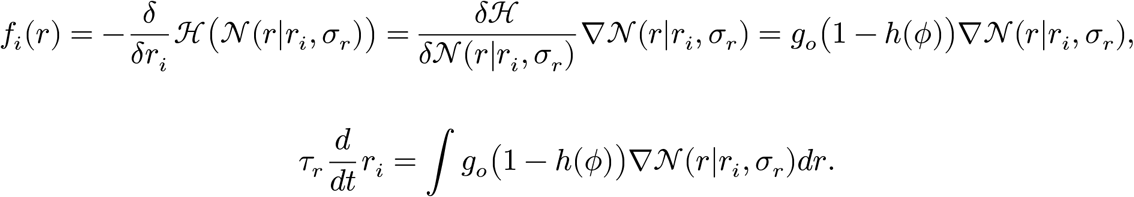

The repulsive interaction is restricted to regions in close proximity to the membrane interface. The term (1 − *h*(*ϕ*)) ensures that the force vanishes in the cell interior and increases as the particle approaches the membrane. Parameters are summarized in **Table 2**.

**Table 2:** Phase-field model parameters.

| Parameter | Description | Value |
| --- | --- | --- |
| Membrane |  |  |
| $\tau$ | Viscous friction coefficient [pN/$\mu m^2$] | 1.0 |
| $\sigma$ | Spatial scale of F-actin [$\mu m$] | 0.6 |
| $V_0$ | Cell area [$\mu m^2$] | 600.00 |
| $M_V$ | Area conservation constraint [$\text{pN}/\mu\text{m}^3$] | 0.1 |
| $g_0$ | Protrusion force [$\text{pN}/\mu\text{m}$ (conc.)<sup>-1</sup>] | 2.0 |
| $\gamma$ | Surface tension [$\text{pN}$] | 2.0 |
| $\epsilon$ | Spatial scale of the phase boundary [$\mu\text{m}$] | 2.0 |
| $\tau_r$ | Viscous friction coefficient [$\text{pN}/\mu\text{m}^2$] | 10.0 |

## Acknowledgements

This study was supported in part by the Moonshot R&D–MILLENNIA Program [grant number JPMJMS2024-9 to H.N.] by Japan Science and Technology Agency (JST), Grant-in-Aid for Transformative Research Areas (B) [grant number 21H05170 to H.N.], Grant-in-Aid for Scientific Research (B) [grant number 21H03541 to H.N.] and JSPS KAKENHI [25H01364 and 25K07242 to N.S.] from the Japan Society for the Promotion of Science (JSPS), Cooperative Study Program of Exploratory Research Center on Life and Living Systems (ExCELLS) [program number 19-102 to H.N.], and Joint Research of the Exploratory Research Center on Life and Living Systems (ExCELLS) [ExCELLS program No. 25EX603].

## Code Availability

Simulations are mainly implemented in C and performed under Python 3.11.0, and the code will be distributed through GitHub after publication.is distributed through GitHub.

## Author Contributions

H.N. conceived the project. M.F., N.S., and H.N. developed the model. M.F. implemented the model and analyzed the data. M.F., N.S., and H.N. wrote the manuscript.

